# Hyperactivity is linked to elevated cortisol levels: comprehensive behavioral analysis in the prenatal valproic acid-induced marmoset model of autism

**DOI:** 10.1101/2025.10.09.676449

**Authors:** Madoka Nakamura, Toru Nakamura, Akiko Nakagami, Keiko Nakagaki, Nobuyuki Kawai, Noritaka Ichinohe

**Affiliations:** Department of Ultrastructure Research, National Institute of Neuroscience, National Center of Neurology and Psychiatry, 4-1-1 Ogawahigasi-cho, Kodaira, Tokyo, 187-8551 Japan; United Graduate School of Agricultural Science, Tokyo University of Agriculture and Technology, 3-8-1 Harumi-cho, Fuchu, Tokyo, 183-8538 Japan; Institute for Datability Science, Osaka University, Techno-alliance bldg. C503, 2-8, Yamadaoka, Suita, Osaka, 565-0871 Japan; Department of Psychology, Japan Women’s University, 2-8-1 Mejirodai, Bunkyo-ku, Tokyo, 112-0015 Japan; Graduate School of Informatics, Nagoya University, Furocho, Nagoya, Aichi, 464-8601 Japan; Academy of Emerging Sciences, Chubu University, 1200 Matsumoto-cho, Kasugai, Aichi, 487-7501 Japan

## Abstract

Hyperactivity is frequently observed in individuals with autism spectrum disorder (ASD) and significantly affects various aspects of life. This underscores the critical need for effective intervention methods tailored to the needs of individuals with ASD. Non-human primate models offer a promising avenue for elucidating the intricate interplay between ASD characteristics and developing individualized therapeutic strategies. This study examined the activity levels and behavioral dynamics in a prenatal valproic acid-induced (VPA) common marmoset model of ASD using ultraminiature data loggers, employing a more detailed approach to behavioral pattern analysis than is traditionally utilized. Although the overall activity levels showed no significant differences, the VPA group exhibited increased activity during specific hours, which is consistent with human ASD studies. Sample Entropy, a statistical measure used to quantify the regularity and unpredictability of time-series data, was higher during daytime in the VPA group, indicating reduced regularity in activity patterns akin to impulsive behavior in ASD. Subtle patterns that were not discernible through simple group comparisons were identified, highlighting the potential of this method as a valuable tool for the behavioral analysis of human ASD. Associations between erratic activity patterns, brief resting intervals, and elevated cortisol levels were observed, all of which correspond to stress phenotypes in individuals with ASD. The findings revealed variations in activity among the adult VPA groups, potentially linked to stress responses. Additionally, VPA juvenile marmosets showed increased locomotor activity in the social interaction test, complementing the adult behavioral findings and suggesting age-dependent manifestations of hyperactivity in this model. This non-human primate model effectively replicates real-world scenarios encountered by individuals with ASD exhibiting hyperactivity, thus holding significant implications for the advancement of personalized therapeutic strategies.

## Introduction

Individuals with autism spectrum disorder (ASD) commonly exhibit a diverse range of complex neurodevelopmental conditions, characterized by difficulties in social interaction, communication, stereotyped behavior, and restricted interests [1,2]. Although hyperactivity is not a core diagnostic feature of ASD under DSM-5/DSM-5-TR [1,3], heightened activity is frequently observed in subgroups of individuals with ASD and is often associated with co-occurring attention-deficit/hyperactivity disorder (ADHD) [4– 7]. In ASD-ADHD co-occurrence, large-scale neuroanatomical deviations— characterized by broadly increased cortical thickness with partly reduced surface area— have been described [8], and externalizing problems such as hyperactivity/impulsivity and attentional difficulties tend to be more severe and pervasive than in ASD alone [9]. Together, these observations underscore the importance of clarifying how activity-related phenotypes intersect with social communication differences in ASD and of developing measurement approaches that respect this heterogeneity.

Within the context of ASD, hyperactivity transcends merely heightened activity levels and is frequently evaluated using tools such as the Behavior Assessment System for Children [10–12]. This instrument relies on subjective reports, primarily from parents and teachers, and captures behavioral indices such as ‘Acts without thinking,’ ‘Is overly active,’ and ‘Interrupts others when they are speaking,’ frequently marked as ‘Often.’ While indicative of hyperactivity, these items also encompass compulsivity and the inability to appropriately modulate behavior, suggesting that hyperactivity in ASD involves broader regulatory challenges rather than the singular symptomatology of excessive movement.

Understanding hyperactivity in ASD presents challenges, including elucidating its mechanisms, interactions with other ASD symptoms, and developing interventions. Recent studies have advanced our understanding of diurnal activity patterns in ASD using actigraphy and wearable devices [13–18]. Actigraphy enables activity monitoring in free-living environments and provides access to raw signals for extracting various physical activity characteristics [19,20]. Although traditional summary variables are prevalent, recent developments in activity pattern analysis involve three distinct methods: analyzing the activity intensity distribution, characterizing the activity duration spread over time, and examining temporal correlations in the activity data. Studies using actigraphy in ASD have revealed distinct activity phenotypes associated with negative skewness shifts linked to acoustic hyper-reactivity, sensory gating alterations, and social and attentional challenges [14].

Since hyperactivity can significantly affect the daily life of individuals with ASD, it is crucial to consider the role of stress in contributing to or exacerbating these behavioral manifestations. Hyperactivity may cause or at least correlate with stress. In fact, hyperactivity involves the dysfunction of the prefrontal cortex [21,22], which is the brain area affected by stress [23]. Moreover, stress is becoming increasingly common among individuals with ASD [24–26]. Individuals with ASD are believed to experience higher levels of stress than their neurotypical peers because of the challenges in coping with change, anticipation, sensory stimuli, and unpleasant daily events [24]. Findings from survey measures revealed that adults with ASD reported significantly greater perceived stress and more stressful life events than typical volunteers, indicating that stress is a clinically significant factor that leads to distress in individuals with ASD, and thus warrants intervention [27].

Animal models hold promise for unraveling the intricate interplay of ASD symptoms and can be instrumental in the development of personalized therapeutic strategies. The intricate and multifaceted nature of ASD presents challenges, prompting scientists to use animal models to replicate crucial ASD aspects. Although mouse models have laid the foundation for the development of animal models of ASD, there are limitations to relying on a single species to study complex human brain disorders [28]. Common marmosets (*Callithrix jacchus*), small New World monkeys, offer distinct advantages over commonly used rodent models for ASD [29,30] because of their close evolutionary proximity to humans and rich social behaviors [31]. Unlike rodents, which are nocturnal and exhibit fragmented sleep patterns even during the day, marmosets are diurnal and closely mirror human activity cycles. Their daytime activities, along with their complex social behaviors and cognitive skills, make them a more suitable and relevant model for studying human-like activity patterns and neurobehavioral processes. This study focused on a marmoset model exposed to valproic acid (VPA) *in utero*, a drug commonly used for its antiepileptic and mood-stabilizing properties, which also acts as a histone deacetylase inhibitor. Notably, VPA exposure during early pregnancy in humans is associated with an increased risk of ASD [32]. These marmosets, ranging from adults to juveniles, not only exhibit ASD-associated behaviors and biological features [33] but also more accurately replicate the human idiopathic ASD brain transcriptome compared to rodent models [34], enhancing our understanding of ASD neurobiology across developmental stages.

This study aimed to capture the associations between hyperactivity, cortisol levels, and behavioral dynamics in adult marmosets with ASD through detailed analyses using actigraphy. By unraveling the interplay between these variables, we sought to enhance our understanding of ASD behavioral phenotypes and contribute to innovative diagnostic and interventional approaches. Home cage activity was recorded in adult VPA-exposed marmosets using actigraphy, employing advanced methodologies, such as the cosinor method [35], behavioral organization analysis [36,37], and Sample Entropy [38,39]. Additionally, this study investigated the correlation between these indices and stress levels, as indicated by salivary cortisol levels. Given the age- and context-dependent variability of hyperactivity in patients with ASD, the relationship between activity and social preference was also explored in juvenile marmosets using a three-chamber test in the presence of unfamiliar conspecifics.

## Methods

### Subjects

This study included 20 common marmosets: 10 adult males and 10 juveniles (Table 1), all bred and housed at the National Center of Neurology and Psychiatry, Tokyo, Japan. In each age cohort, we included five prenatal valproic acid-exposed (VPA) ASD-model animals and five unexposed (UE) controls. No formal power calculation was performed. The sample size was determined based on prior publications using the same VPA-exposed marmoset model and the practical limitations of non-human primate gestational studies. No randomization was used for group allocation or experimental ordering because prenatal VPA exposure predetermined the model group. Control animals were age- and sex-matched and housed under identical conditions. Among the ten adult marmosets involved in the physical activity analysis (Fig. 1A), one VPA individual (animal ID: 16017; Table 1) was excluded from the cortisol experiment because of an inability to be trained in saliva collection method which involves chewing cotton swabs prior to the commencement of collection. The exclusion criterion (failure in training or incomplete behavioral data) was pre-defined before the analysis. The adult marmosets selected for the physical activity monitoring (Fig. 1A) and cortisol level measurement exhibited an age spectrum ranging from 2.0 to 4.8 years (mean ± SEM = 3.6 ± 0.3 years). Meanwhile, the juvenile marmosets who participated in the three-chamber test (Fig. 1B) were aged in the range of 15 to 19 weeks (mean ± SEM = 17 ± 0.2 weeks). Each adult marmoset was housed individually in stainless-steel cages located within a shared colony room. In contrast, juvenile marmosets cohabited with their parents and siblings within the same enclosure. A consistent light cycle was maintained, with 12 h of light followed by 12 h of darkness, commencing at 07:00 h. Environmental conditions were rigorously controlled, with temperature and humidity maintained at 28 ± 2 °C and 50 ± 10%, respectively. Marmosets were given food (CMS-1, CLEA Japan) twice daily, in the morning and afternoon, and had unrestricted access to water that was readily available in the front mesh of their cages. No blinding was implemented for investigators or during outcome assessment; all measurements were obtained objectively using quantitative methods. All research procedures were approved by the Animal Experiment Ethics Committee of the National Center of Neurology and Psychiatry and were in accordance with the NIH Guide for the Care and Use of Laboratory Animals.

**Fig. 1:**
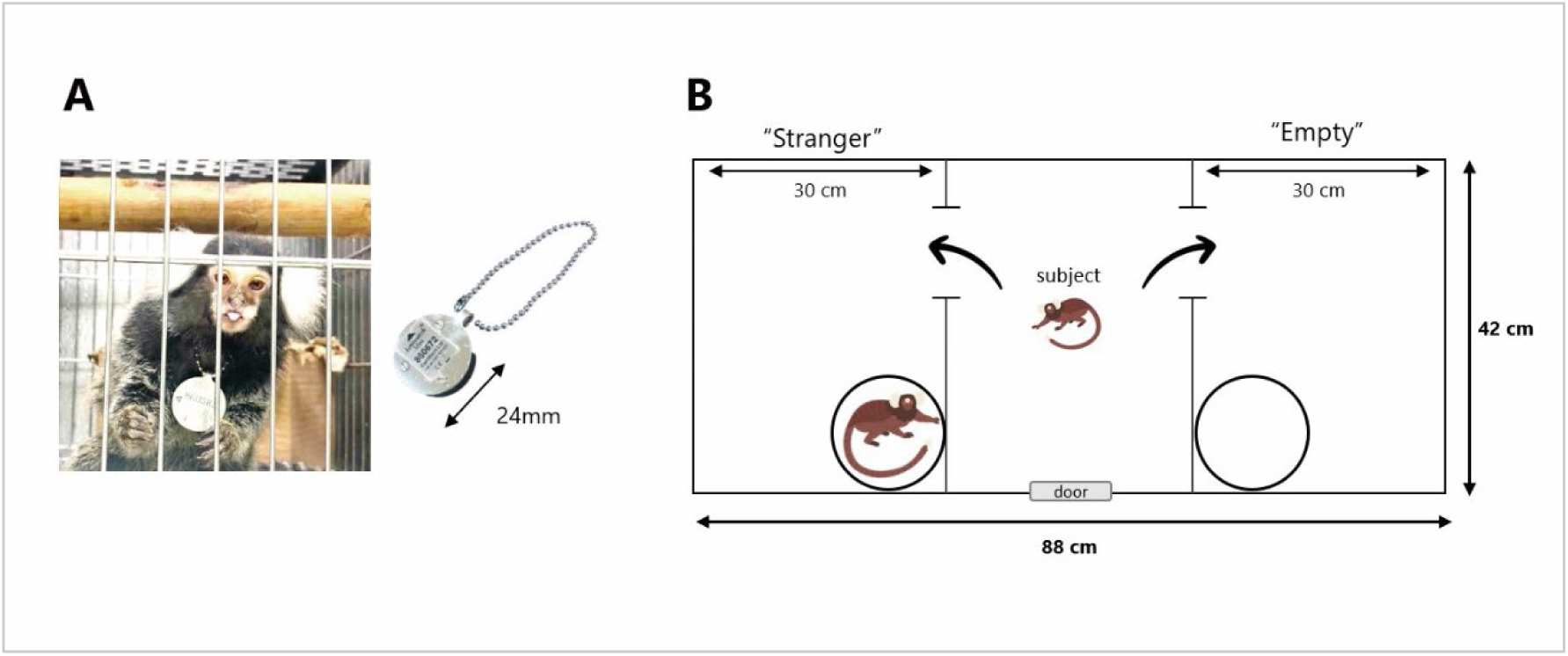
Experimental equipment. **A** Actigraphy-based data logger device “Actiwatch Mini^®^” (*right*) and a photograph of an adult test subject wearing the device (*left*). **B** Three-chamber test apparatus layout for juvenile subjects. The locations of the “stranger” and “empty” areas were counterbalanced. The subject marmoset entered the central chamber voluntarily, and the sliding doors were closed. Both doors of the side chambers were then opened to allow the animals to explore the entire test apparatus. An unfamiliar adult marmoset was placed in one of the side chambers, in a clear plastic tube with air holes on the lid.

**Table 1:** Subject Information.

| *Experiment* | *Animal ID* | *Group* | *Sex* | *Age (week)* | *Weight (g)* |
| --- | --- | --- | --- | --- | --- |
| Activity Analysis | 14030 | VPA | Male | 252 | 416.7 |
|  | 14031 | VPA | Male | 252 | 431.3 |
|  | 16017 | VPA | Male | 161 | 507.0 |
|  | 16043 | VPA | Male | 152 | 410.6 |
|  | 17024 | VPA | Male | 104 | 364.9 |
|  | 14078 | UE | Male | 226 | 382.7 |
|  | 15033 | UE | Male | 200 | 393.1 |
|  | 15034 | UE | Male | 200 | 397.6 |
|  | 15050 | UE | Male | 187 | 422.5 |
|  | 16089 | UE | Male | 139 | 543.5 |
| 3-chamber test | 19061 | VPA | Male | 15 | 167.6 |
|  |  |  |  | 16 | 171.3 |
|  |  |  |  | 17 | 185.2 |
|  |  |  |  | 18 | 190.3 |
|  |  |  |  | 19 | 193.6 |
|  | 19062 | VPA | Female | 15 | 184.2 |
|  |  |  |  | 16 | 187.8 |
|  |  |  |  | 17 | 200.8 |
|  |  |  |  | 18 | 211.2 |
|  |  |  |  | 19 | 217.3 |
|  | 19072 | VPA | Male | 15 | 163.2 |
|  |  |  |  | 16 | 170.0 |
|  |  |  |  | 17 | 186.7 |
|  |  |  |  | 18 | 197.7 |
|  |  |  |  | 19 | 208.5 |
|  | 19143 | VPA | Female | 15 | 187.5 |
|  |  |  |  | 16 | 196.0 |
|  |  |  |  | 17 | 199.0 |
|  |  |  |  | 18 | 211.0 |
|  |  |  |  | 19 | 209.7 |
|  | 19145 | VPA | Female | 15 | 152.3 |
|  |  |  |  | 16 | 157.6 |
|  |  |  |  | 17 | 160.5 |
|  |  |  |  | 18 | 168.8 |
|  |  |  |  | 19 | 173.5 |
|  | 19091 | UE | Female | 15 | 172.2 |
|  |  |  |  | 16 | 184.2 |
|  |  |  |  | 17 | 199.8 |
|  |  |  |  | 18 | 198.6 |
|  |  |  |  | 19 | 205.0 |
|  | 19092 | UE | Male | 15 | 170.9 |
|  |  |  |  | 16 | 191.0 |
|  |  |  |  | 17 | 202.4 |
|  |  |  |  | 18 | 207.1 |
|  |  |  |  | 19 | 208.3 |
|  | 19093 | UE | Male | 15 | 177.7 |
|  |  |  |  | 16 | 195.6 |
|  |  |  |  | 17 | 210.9 |
|  |  |  |  | 18 | 214.8 |
|  |  |  |  | 19 | 209.8 |
|  | 19099 | UE | Female | 15 | 202.7 |
|  |  |  |  | 16 | 207.9 |
|  |  |  |  | 17 | 215.1 |
|  |  |  |  | 18 | 222.3 |
|  |  |  |  | 19 | 228.1 |
|  | 19100 | UE | Female | 15 | 187.8 |
|  |  |  |  | 16 | 200.8 |
|  |  |  |  | 17 | 201.6 |
|  |  |  |  | 18 | 206.9 |
|  |  |  |  | 19 | 222.1 |

### Valproic acid treatment during pregnancy

The VPA model of ASD in common marmosets was established as described previously [33,34,40]. Briefly, 4% VPA solution was prepared in 10% glucose solution and administered orally to the marmosets during pregnancy. The dams in the VPA group received a daily dose of 200 mg/kg. The VPA administration regimen was maintained from days 60 to 66 after conception, with a total of seven treatments. This specific timeframe was chosen to align with the analogous administration period (E12) used in the development of VPA-exposed rodent models of ASD. Both VPA-exposed and unexposed (UE) marmosets were housed in separate cages. To determine the precise timing of pregnancy, blood progesterone levels were regularly monitored in VPA-exposed dams, following the same protocol used for UE dams. Notably, all VPA-exposed dams exhibited tolerance to the medication without any instances of vomiting, and displayed no signs of abnormal pregnancy or delivery. In contrast, UE marmoset dams did not receive VPA during this critical period. The offspring of VPA-exposed marmosets did not manifest any malformations or discrepancies in body weight when compared to their UE subjects and were used as the VPA marmoset group in this study.

### Actiwatch Mini®

Actiwatch-Mini® (Cambridge Neurotechnology Ltd., UK) is a non-invasive ultraminiature actigraphy-based data logger device, specially developed for monitoring activity in animals (Fig. 1A). It is based on the piezoelectric accelerometer technology, which records the intensity, amount, and duration of movements in all directions [41–43]. This activity acquisition system is based on miniaturized accelerometer technologies that are currently used for human activity monitoring, but have also been tested for activity monitoring in small non-human mammals. To record locomotor activity, an Actiwatch-Mini® was placed by means of a neck collar that was accepted by the animal without any apparent disturbance for the average of 22 days (mean ± SEM = 22.2 ± 1.1 days). All subjects were first habituated to an aluminum ball-chain collar for one week prior to data collection. Total activity was recorded as the result of all movements, including different behaviors, such as drinking, feeding, walking, and grooming, as well as all conscious and unconscious movements independent of the animal’s position, such as lying or standing. Activity was monitored at a sampling interval of 1 min. During recording, the activity counts were stored in the Actiwatch as one-minute epochs. All data were exported to a PC via a USB coupled reader for analysis using Actiwatch® Activity & Sleep Analysis 7 software.

### Physical activity analysis

#### Cosinor analysis

Sleep-wake biological rhythmicity was evaluated using a traditional cosinor-based analysis, which fits a linear combination of sinusoidal waveforms with specific periods. A single-component cosinor with a 24 h period is a fundamental approach commonly used to infer endogenous circadian rhythms embedded in biological signals. This single cosinor method fits the equation *y*(*t*) = *M* + *Acos*(ω_24_*t* + ф) to the time-series data *y*(*t*), where *M* denotes the mesor (a rhythm-adjusted mean), *A* is the amplitude of the 24 h period (the difference between the mean and peak), ф is the acrophase (the time of the peak of the fitted rhythm) of the 24 h periodicity (2π/ω_24_), and *t* represents time.

Physical activity, known to encompass a significant 12 h harmonic component indicative of a semi-circadian rhythm, cannot be sufficiently characterized using the traditional single-cosinor method in terms of rhythmicity. To address this limitation, double cosinor analysis was used, which involves the simultaneous fitting of two cosine functions: one corresponding to a 24 h period (2π/ω_24_) and the other to 12 h period (2π/ω_12_). The fitting function was formulated as *y*(*t*) = *M* + *A*_24_*cos*(ω_24_*t* + ф_24_) + *A*_12_*cos*(ω_12_*t* + ф_12_), where *M* denotes the mesor of the model, *A*_24_ is the amplitude of the 24 h period, *A*_12_ is the amplitude of the 12 h period, and ф_24_ and ф_12_ are the acrophases of the 24 h and 12 h period, respectively. This analysis was conducted after converting the activity data into aggregated values over 10 min intervals for each subject. The models were fitted to the entire dataset for each individual on a daily basis, followed by the calculation of the average values of the fitting parameters.

#### Behavioral organization analysis

To quantitatively assess the rest-activity properties of the physical activity data, the behavioral organization analysis developed by Nakamura et al. was utilized [36,37]. Based on previous studies, the cumulative probability distribution *P*(*x* ≥ *a*) for durations *a* of both the resting and active periods was estimated by integrating their probability density functions. Resting periods were defined as the durations during which activity counts remained below a predefined threshold, whereas active periods were defined as the durations during which activity counts consistently exceeded this threshold. In our study, the threshold was set as the overall daily mean of the nonzero activity counts to minimize the effects of daily variations attributable to slight differences in the mounting positions of the devices. Both cumulative distributions were characterized by fitting the active durations with a stretched exponential functional form *P*(*x* ≥ *a*) = exp (−α^β^), and the resting durations with a power-law form *P*(*x* ≥ α) = Aα^−γ^. This method provided estimates for the fitting parameters α and β in active periods and the scaling exponent γ in resting periods. The fitting range spanned from 10 to 100 min for the cumulative distributions of active periods, and α ≥ 3 min for resting periods.

#### Sample Entropy

Sample Entropy is a measure of complexity used to evaluate the irregularity or unpredictability of time-series data. It has been extensively applied to analyze the complexity of various signals, including heart rate variability [38,39], electroencephalogram data [44], and behavioral data [45]. For this analysis, data collected during daytime were used to focus on diurnal activity and minimize the influence of nocturnal rest. Sample Entropy calculates the negative natural logarithm of the conditional probability that a dataset of length *N* that has repeated itself for *m* points within a certain tolerance *r* will continue to do so for *m*+1 points, excluding identical matches. Detailed descriptions of the algorithm are available in the literature [38,39]. To ensure consistency in the analysis, activity data were normalized to have a zero mean and a standard deviation of 1. Following prior research, *m* = 2 and *r* = 0.2 were selected for the calculations.

### Salivary collection and assay

To determine the correlation between activity levels and stress-related hormone levels within an individual, the salivary collection methods and cortisol samples described in our previous study were used [46]. Subjects were not captured by the experimenter, and saliva was collected under free-ranging conditions in their home cages. Saliva samples were collected once a month from each individual, and the average of three samples was used for correlation analysis of the individual’s cortisol level. Cortisol levels (μg/dL) were measured using an AIA-360 Automated Immunoassay Analyzer with AIA-pack cortisol test cups (Tosoh Corporation, Tokyo, Japan) and averaged per subject across all saliva collections either in the morning or evening.

### Three-chamber test

Five juvenile marmosets (*n* = 2 males and *n* = 3 females) that underwent early developmental exposure to VPA and another group of five juvenile marmosets (*n* = 2 males and *n* = 3 females) that remained unexposed to VPA (UE group) were selected for the three-chamber test. The assessments, conducted between the ages of 15 and 19 weeks to align with the standard diagnostic period for ASD in humans [47], involved each marmoset participating in a three-chamber test once a week for five trials per individual. An unfamiliar adult marmoset (*n* = 11 males and *n* = 12 females) was placed in one of the side chambers and met target juvenile animals of the same sex only once. These unfamiliar adults were selected because they were not kin to the subjects and had no previous interactions with them that might have provided visual, auditory, or olfactory cues. This study employed identical experimental methodologies and apparatus (Fig. 1B) as described in our previous publication [33]. The time spent in the chambers and the number of transitions among the chambers were measured using Observer XT 11 software (Noldus Information Technology, Netherlands).

### Statistical analysis

All values are expressed as the mean ± standard deviation to indicate within-group variation, and p-values < 0.05 were considered statistically significant. A two-tailed Student’s *t-*test was used to compare physical activity parameters across groups when data distributions were considered approximately normal and variances were similar between groups. A two-factor repeated-measures analysis of variance (ANOVA) with a Bonferroni post-hoc test was used to assess the hourly changes in physical activity levels. Analyses were performed using SAS version 3.81 (SAS Institute, Cary, NC, USA). For the three-chamber test data, two-factor repeated-measures ANOVA and Student’s *t-*test were performed using JMP version 17.2.0 (SAS Institute, Cary, NC, USA).

## Results

### Home-cage activity analysis in adult marmosets

The activity levels and behavioral dynamics of the VPA and UE groups were examined using ultraminiature data loggers, with an analysis of behavioral patterns that were more intricate than those typically employed. The experimental setup depicted in Fig. 1A was used to evaluate the activity levels within the respective home cages. Activity levels over the entire measurement period revealed an enhanced propensity in the VPA-induced marmosets compared to the UE subjects (Fig. 2A). The average daily activity in the VPA group showed an increasing tendency, but was not statistically significant (*p* = 0.1603, Student’s *t*-test; Fig. 2B), whereas there were no significant differences in nighttime activities (7 p.m. to 7 a.m.; *p* = 0.8152, Student’s *t*-test; Fig. 2C). However, substantial group differences were observed in the patterns of hourly physical activity levels. Both the groups showed a bimodal activity pattern during the diurnal period: an increase from 7:00 a.m. (T7) when the rearing room electricity was activated, followed by a decline at approximately 1:00 p.m. (T13), and a peak at approximately 7:00 p.m. (T19) when the electricity was deactivated (Fig. 2D). Hourly physical activity increased in the VPA group during the morning (T8-T10; 8 a.m.-10 a.m.), as well as from 7:00 p.m. to 8:00 p.m. (T19-T20), compared to the UE group. There was no statistically significant difference in the weight of the subjects measured immediately after the experiment between the two groups (Fig. S1).

**Fig. 2:**
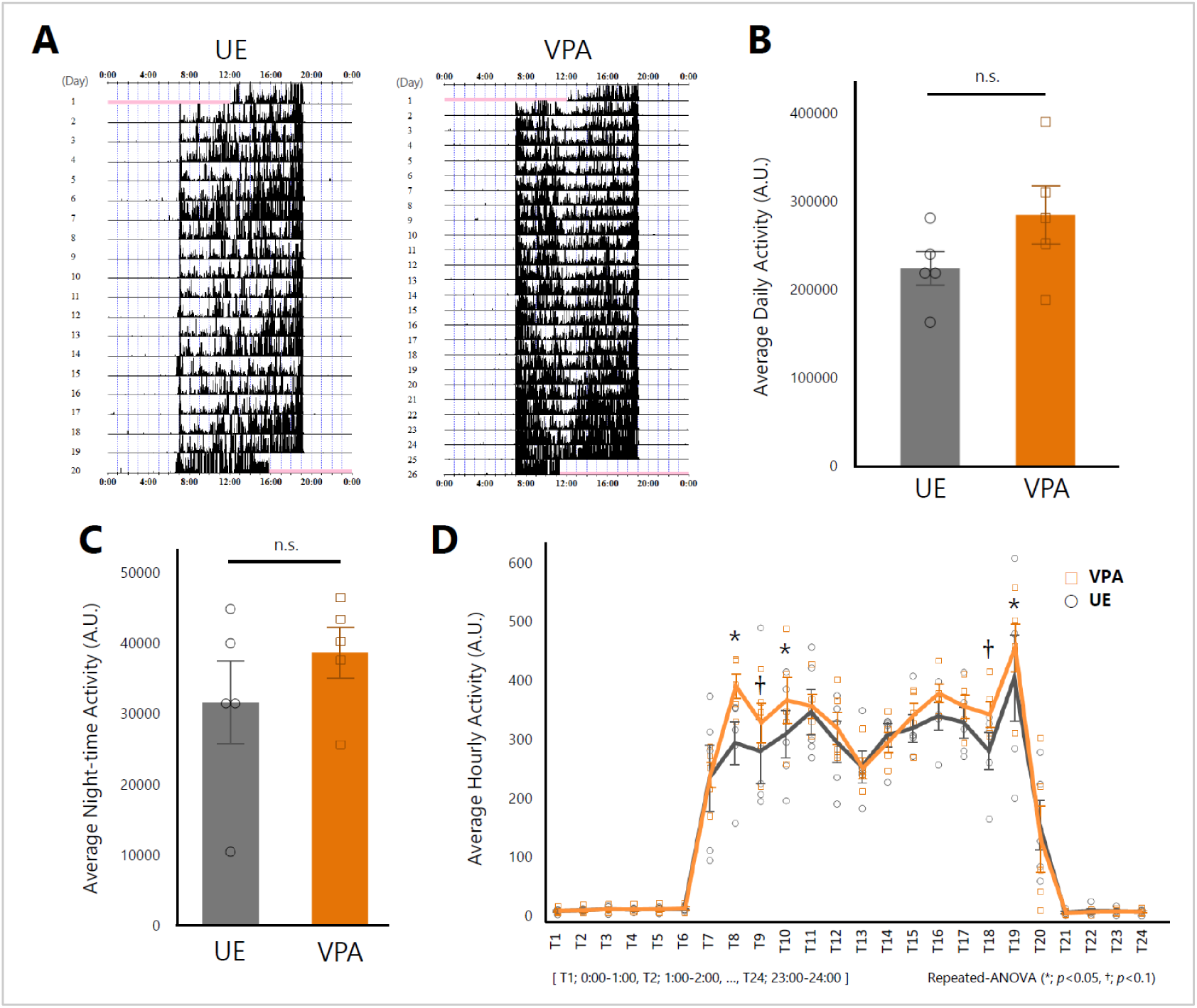
Home cage activity analysis in the valproic acid-exposed and unexposed groups. **A** Representative activity levels over the entire measurement period in valproic acid-exposed (*right*) and unexposed marmosets (*left*). **B** Average daily activity in the UE and VPA groups (*n* = 5 per group). *p* = 0.1603 for two-tailed Student’s *t*-test. **C** Average night-time activity in the UE and VPA groups. *p* = 0.8345 for two-tailed Student’s *t*-test. **D** Average hourly activity levels in the VPA and UE groups. Values are presented as mean ± standard deviation. \**p* < 0.05, †*p* < 0.1 for repeated ANOVA.

The levels and timing of the circadian activity rhythms were estimated using the cosinor analysis method. Cosinor analysis, utilizing both single and double cosinor methods, revealed no significant differences in the fitting parameters between the VPA and UE groups (Table 2; Fig. 3A). This indicates that there were no significant group differences in the magnitude of peak activity levels and timing (acrophase).

**Fig. 3:**
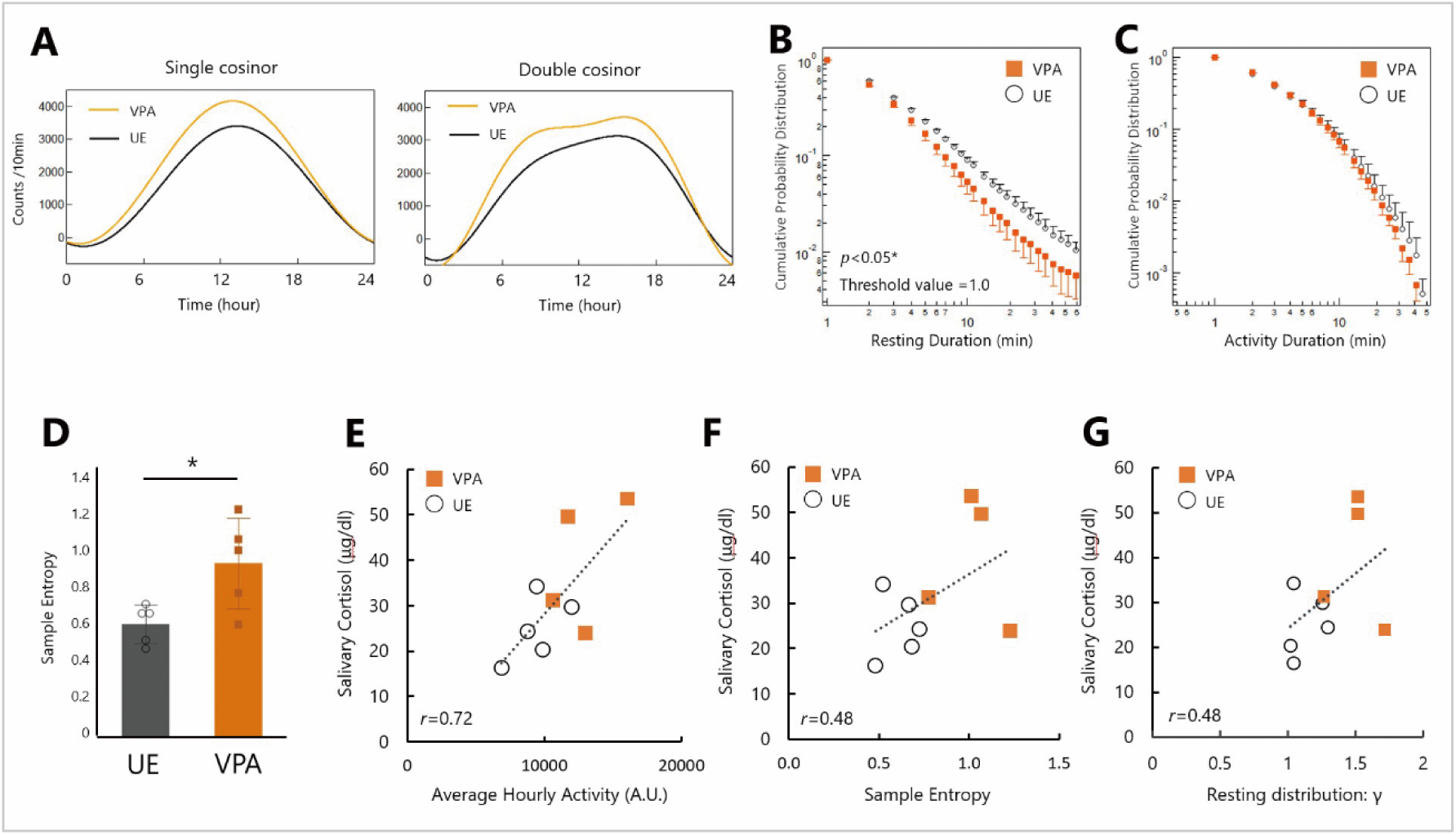
Physical activity analysis in the valproic acid-exposed and unexposed groups. **A** Cosinor curves for single (*left*) and double (*right*) cycles depicting the mean fitting parameter values for each group (*n* = 5 for each group). **B** Cumulative probability distributions for the resting periods in both the groups. **C** Cumulative probability distributions for active periods in both the groups. **D** Average sample entropy in the UE and VPA groups. Values are presented as mean ± standard deviation. \**p* = 0.0269 for two-tailed Student’s *t-*test. Scatter plot (*n* = 5 for UE, *n* = 4 for VPA) showing the relationship between salivary cortisol levels and **E** the average hourly activity levels [*F*(1,7) = 7.6770, *r* = 0.72, *p* = 0.0277], **F** sample entropy [*F*(1,7) = 2.1297, *r* = 0.48, *p* = 0.1878], and **G** resting distribution parameters γ [*F*(1,7) = 2.1524, *r* = 0.48, *p* = 0.1858].

**Table 2:** Comparison of physical activity parameters among groups.

| Parameters | UE | VPA | $p$-value |
| --- | --- | --- | --- |
| ***Basic statistics (with nocturnal data)*** |  |  |  |
| Mean | 157.2 ± 29.7 | 199.0 ± 51.6 | $p = 0.16$ |
| SD | 317.1 ± 44.4 | 352.0 ± 54.3 | $p = 0.30$ |
| Skewness | 2.8 ± 0.7 | 2.4 ± 0.8 | $p = 0.37$ |
| Kurtosis | 10.2 ± 5.9 | 8.0 ± 7.2 | $p = 0.60$ |
| ***Basic statistics (without nocturnal data)*** |  |  |  |
| Mean | 299.2 ± 54.4 | 383.1 ± 100.4 | $p = 0.14$ |
| SD | 385.0 ± 61.7 | 409.7 ± 58.7 | $p = 0.54$ |
| Skewness | 1.8 ± 0.5 | 1.5 ± 0.7 | $p = 0.38$ |
| Kurtosis | 4.7 ± 3.7 | 3.7 ± 4.6 | $p = 0.70$ |
| **Single cosinor method** |  |  |  |
| Mesor: $M$ | 1569.3 ± 310.3 | 1991.0 ± 525.3 | $p = 0.16$ |
| Amplitude (24h): $A$ | 1875.2 ± 337.0 | 2207.0 ± 557.7 | $p = 0.29$ |
| Acrophase (24h) [rad]: $\varphi$ | 2.86 ± 0.32 | 2.88 ± 0.08 | $p = 0.90$ |
| **Double cosinor method** |  |  |  |
| Mesor: $M$ | 1569.3 ± 310.3 | 1991.6 ± 526.4 | $p = 0.16$ |
| Amplitude (24h): $A_{24}$ | 1875.2 ± 337.0 | 2206.3 ± 556.6 | $p = 0.29$ |
| Acrophase (24h) [rad] $\varphi_{24}$ | 2.86 ± 0.32 | 2.88 ± 0.08 | $p = 0.90$ |
| Amplitude (12h): $A_{12}$ | 550.2 ± 98.7 | 751.9 ± 257.3 | $p = 0.14$ |
| Acrophase (12h) [rad]: $\varphi_{12}$ | 3.04 ± 0.51 | 2.84 ± 0.53 | $p = 0.56$ |
| **Behavioral organization analysis** |  |  |  |
| Resting distribution: $\gamma$ | 1.10 ± 0.1 | 1.38 ± 0.22 | $p < 0.05^*$ |
| Active distribution: $\alpha$ | 0.86 ± 0.28 | 0.67 ± 0.22 | $p = 0.25$ |
| Active distribution: $\beta$ | 0.60 ± 0.13 | 0.67 ± 0.12 | $p = 0.42$ |
| **Sample entropy** | 0.61 ± 0.11 | 0.94 ± 0.24 | $p < 0.05^*$ |

Values are represented as mean ± standard deviation.

Behavioral organization analysis revealed characteristic rest-active patterns in the behavioral dynamics (Fig. 3B and 3C). The distribution of resting periods in the VPA group was reduced compared with that in the UE group (Fig. 3B), particularly at longer durations, implying less persistence of resting periods in the VPA group. The resting period distributions in both the groups followed a power-law distribution over one decade (2 min–30 min) with significantly different scaling exponents of γ = 1.38 ± 0.22 and γ = 1.10 ± 0.1 for the VPA and UE groups, respectively (*p* < 0.05; see Table 2). This difference suggests that the VPA group tended to reactivate more quickly once they entered the resting-state, resulting in shorter resting periods. On the other hand, the distribution of active periods in both the groups had a stretched exponential form (Fig. 3C) with no significant difference in the fitting parameters α and β (*p* = 0.25 and *p* = 0.42, respectively; see Table 2).

A group comparison of the Sample Entropy values evaluated from diurnal physical activities is presented in Fig. 3D. The mean Sample Entropy of the VPA group was significantly higher than that of the UE group, indicating increased irregularity or unpredictability of behavioral dynamics in the VPA group (*p* = 0.027, Student’s *t*-test; Fig. 3D). Interestingly, a significant positive correlation was found between the average hourly activity and salivary cortisol levels upon waking collected during the non-Actiwatch-wearing period (*r* = 0.72, *p* = 0.0277; Fig. 3E). In addition, a moderate positive correlation was observed between Sample Entropy and salivary cortisol levels (*r* = 0.48, *p* = 0.1878; Fig. 3F), as well as between the resting period distribution parameter γ and salivary cortisol levels (*r* = 0.48, *p* = 0.1858; Fig. 3G).

### Three-chamber test analysis in juvenile marmosets

Adult home-cage analyses showed no group difference in daily means but modest, time-specific increases in activity in the VPA group. To examine age- and context-dependent expression, we tested juveniles under parental care in a three-chamber social-preference test. Juvenile marmosets underwent the three-chamber test five times between 15 and 19 weeks of age. All subjects were able to move freely in the “stranger” area with a strange same-sex adult marmoset in the container, the “empty” area with an empty container, and the central area (Fig. 1B). The UE group spent significantly longer time in the “stranger” area compared to the “empty” area, whereas no significant differences were observed between the chambers in the VPA group (*p* < 0.0001, two-way repeated-measures ANOVA, Fig. 4A). The social preference index was examined to accurately quantify alterations in sociability. It was calculated as follows: (time (%) spent in the stranger chamber - time (%) spent in the empty chamber)/100, with a value closer to 1 indicating a greater tendency to be in the stranger area. The social preference index of the VPA group was much lower than that of the UE group (*p* = 0.0426, Student’s *t*-test; Fig. 4B). When the number of times the subjects moved into the three compartments during the experimental period was counted, the VPA group moved significantly more than the UE group (*p* = 0.0092, Student’s *t-*test; Fig. 4C). There was a significant negative correlation between the number of times a parcel was moved and the percentage of time spent in the “stranger” area (*r* = −0.71, *p* = 0.0212, Fig. 4D).

**Fig. 4:**
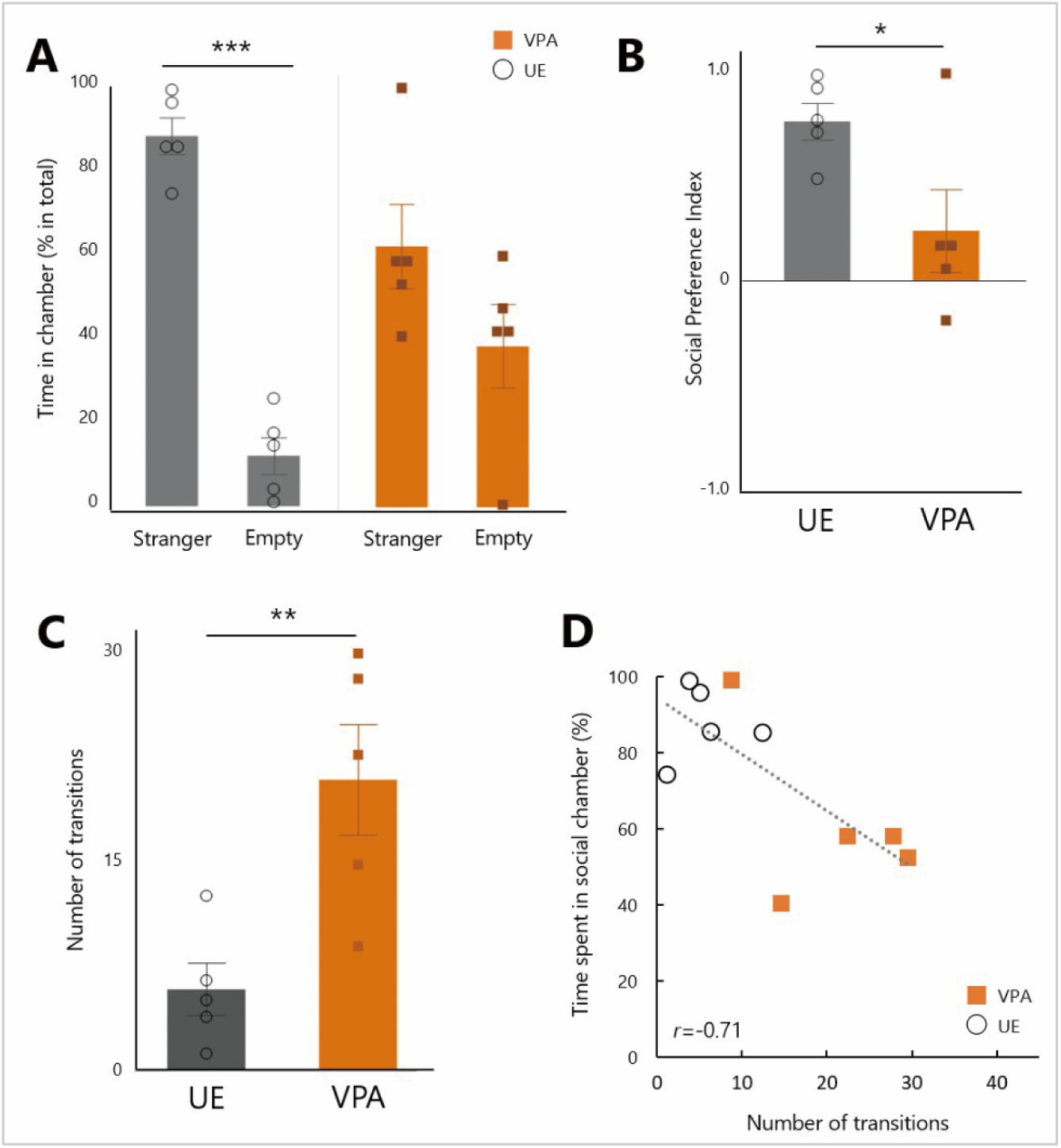
Three-chamber test analysis data in juvenile subjects. **A** The time (% in total) spent in the stranger and empty chambers. UE marmosets spent significantly more time in the stranger area (\*\*\**p* < 0.0001, two-way repeated-measures ANOVA followed by Tukey’s HSD post-hoc test; *n* = 5 for each group). **B** Social preference index in the UE and VPA groups (*n* = 5 per group; \**p* = 0.0426, two-tailed Student’s *t*-test). **C** Number of transitions among chambers in the UE and VPA groups (\*\**p* = 0.0092, two-tailed Student’s *t*-test). **D** Scatter plot showing the relationship between the number of transitions and time (%) spent in the social (stranger) chamber. *F*(1,7) = 8.1627, *r* = −0.711, *p* = 0.0212.

## Discussion

In this study, Actiwatch® technology was used to closely examine the activity patterns of adult VPA-induced ASD marmosets in their home cages, employing a more intricate approach to behavioral pattern analysis than the traditional method. In addition, the potential links between these activity patterns and stress were investigated by measuring cortisol levels, a well-established stress marker. The findings suggest that ASD model marmosets exhibit variations in activity that may be linked to stress responses, as indicated by their cortisol profiles. In addition, the activity levels of juvenile marmosets exposed to VPA were explored in unfamiliar social environments and associations between their activity levels and social interactions were observed.

The activity analyses showed increases during morning and evening hours in the ASD model, despite no significant differences in the overall mean activity levels between the VPA and control groups. This finding aligns with recent human ASD studies utilizing actigraphy analysis [14,15,18], where detailed scrutiny revealed heightened activity intricacies rather than broad activity measures. Recent studies have indicated that simple measures, such as mean activity levels, do not capture the variability or complexity of activity patterns in neurodevelopmental disorders [48,49]. The subtle differences observed in the overall activity patterns of VPA-exposed marmosets and human studies [14–16,18] may arise from various factors, including device specifications, measurement environments, and the heterogeneity of ASD symptoms. These factors are essential for translating findings from marmosets to humans.

A recent study revealed a discrepancy in behavior among four representative mouse models of ASD, which exhibited a burst of hyperactivity in a limited-time open-field test in an unfamiliar environment, followed by significant hypoactivity during prolonged recordings in the familiarity of a home cage [50]. Notably, Bass et al. observed an increased activity in VPA-exposed mice when introduced to a running wheel within their cage, which was not observed in the absence of a running wheel [51]. These findings underscore the significant influence of environmental context on activity levels, highlighting the need for careful data interpretation. Additionally, we observed a shift towards hyperactivity in juvenile ASD marmosets when unfamiliar adults were encountered in a three-chamber setting (Fig. 4C). Although adults and juveniles were assessed with different assays, this juvenile finding still offers valuable insights into the understanding of activity patterns in ASD models and contributes to the knowledge of activity levels in VPA-exposed marmoset offspring in social settings, emphasizing the variability of this phenotype across different environments and developmental stages.

Daily physical activity is a multifaceted behavior that requires consideration of multiple dimensions such as intensity, duration, and complexity. Wearable devices such as Actiwatch® have proven useful in monitoring the physical activity of patients with psychiatric and neurological disorders in free-living environments [52–55]. Time-series data from wearable devices offer the opportunity to extract various features of daily activities, leading to the proposal of various analytical methods [17]. In the present study, Sample Entropy, a statistical measure used to quantify the regularity and unpredictability of time-series data, was used to assess the predictability of activity in VPA-exposed marmosets (Fig. 3D). This provided a statistical evaluation of the regularity and unpredictability of the activity patterns. The VPA-induced ASD marmosets exhibited higher entropy during the daytime, indicating a reduction in behavioral predictability, potentially capturing aspects of impulsive and unpredictable behavior often displayed by individuals with ASD [48]. High entropy has been reported in bipolar disorder during manic and mixed states and in patients with schizophrenia, indicating its relevance in various psychiatric conditions [48,56,57]. Linking video recordings with activity data to investigate behaviors that contribute to increased entropy may provide a more accurate characterization of VPA-exposed marmosets and serve as a benchmark for interpreting similar records in individuals with ASD.

Our investigation revealed a connection between erratic and excessive activity patterns with brief resting intervals and elevated cortisol levels. These findings may be related to well-documented adverse effects associated with prolonged steroid use, such as mood fluctuations, anxiety, depression, and excitability. Additionally, the stress phenotype commonly observed in individuals with ASD may contribute to increased cortisol levels and hyperactivity in VPA-exposed marmosets. Research on individuals with ASD has connected hypoactivity and spontaneous hyperactivity to a predisposition towards negative skewness, which is correlated with acoustic hyper-reactivity and deficits in sensory gating [14]. Such associations implicate prefrontal cortex (PFC) dysfunction, as stress is known to weaken prefrontal networks, and hyperactivity has been linked to PFC dysfunction [21,22]. Consistent with this, the electrocorticogram (ECoG) study using this VPA marmoset model reported erroneous predictive coding across brain hierarchies, suggesting PFC-related dysfunction that may underlie the current findings [58]. We therefore hypothesize a multidirectional model in which auditory hypersensitivity increases stress response, leading to elevated cortisol levels. Simultaneously, increased stress may decrease sensory filtering, leading to increased auditory sensitivity. Sensory overload may trigger hyperactivity as a coping mechanism, which further increases arousal and stress. This interplay could sustain a self-perpetuating cycle of hypersensitivity, stress, and hyperactivity, with each component potentially amplifying the others. The persistence of auditory hypersensitivity across the brain hierarchies may serve as a catalyst for this cycle, underlying the chronic and interconnected nature of these symptoms in ASD.

Further studies in children with ASD have identified a correlation between skewed activity patterns and difficulties in attention, hyperactivity, and peer interactions, suggesting a connection between atypical activity patterns and a range of ASD symptoms [13]. A correlation between the vigor of inter-chamber movement and social preference was observed in juvenile marmosets in this study, reinforcing the notion that altered activity patterns may stem from mechanisms linked to a broader ASD phenotype. These observations in both human studies and our marmoset model provide additional support for the proposed multidirectional model, suggesting that the interplay between hypersensitivity, stress, and hyperactivity may have far-reaching effects on social behavior and cognitive functioning in ASD.

This study has potential limitations that could provide guidance for future research. First, the interpretation of our findings is subject to constraints related to the study cohort. The sample size was necessarily limited by ethical and logistical constraints inherent in a non-human primate gestational model, resulting in small cohorts (*n* = 5 per group for adults and juveniles). While this constraint may have contributed to the lack of statistical significance in basic metrics such as overall mean activity levels (*p* = 0.1603; Fig. 2B), the depth of advanced analytical methods employed provided significant insights into behavioral dynamics. Second, the adult cohort utilized for actigraphy and cortisol analysis consisted exclusively of males, meaning the sex distribution was imbalanced. This limitation restricts the generalizability of the findings across sexes and underscores the need for sex-balanced samples in future work. Third, we observed a trend toward a group difference in juvenile body weight (*p* = 0.0878; Fig S2). We also acknowledge substantial age variability in the adult cohort (Fig S1). While these factors were not statistically significant in their group differences, they may contribute to the heterogeneity observed in the behavioral measures. Fourth, juvenile home-cage actigraphy was not performed due to device-mass constraints, and adult three-chamber testing was not performed because of welfare and ethological considerations. In parallel, to accommodate adult testing with maximal attention to animal welfare, we are developing a non-contact, choice-based social preference apparatus that relies on voluntary approach and avoidance without restraint while preserving ethological relevance. For juveniles, given the mass constraint of Actiwatch, we will quantify behavior with a markerless pipeline based on DeepLabCut [59] that requires no attached device. This may allow us to analyze activity levels in juveniles and investigate autism-like behaviors, such as a lack of interaction with parents and other adult individuals. Future studies should examine the potential of this method for measuring activity levels and behaviors under various environmental conditions.

In conclusion, our study unveiled striking parallels in activities and stress responses, as well as their interrelations, thereby positioning marmosets as a crucial model for deciphering the interplay, causality, and underlying mechanisms among ASD phenotypes. The VPA-exposed marmoset, with its diverse and detectable biological and behavioral features [33,34,40,46,60,61], holds promise for unveiling the mechanisms underlying the various symptoms of ASD. We propose that this non-human primate model faithfully mirrors the typical circumstances encountered by individuals with ASD who exhibit hyperactivity, offering potential avenues for refining personalized therapeutic interventions. Investigating the impact of pharmacological, electrical, and genetic interventions, as well as behavioral therapy, on the correlations between the parameters identified in our current and past research on VPA-exposed marmosets could yield deeper insights into symptom-specific mechanisms. These approaches not only aim to enhance our understanding of ASD, but also promote the development of targeted therapeutic strategies tailored to individual neurobiological profiles.

## Data availability statement

The original contributions presented in this study are included in the article/supplementary material. Further inquiries can be directed to the corresponding authors.

## Ethics statement

This animal study was reviewed and approved by the regulations of the National Center of Neurology and Psychiatry (NCNP), Tokyo, Japan.

## Author contributions

MN designed, conducted experiments, analyzed data and wrote the manuscript. TN analyzed data and helped with the manuscript. AN designed and conducted experiments. KN managed the production and physical condition of the animals. NK helped with the design of the studies and offered comments on the manuscript. NI designed, supervised and helped with the manuscript. All authors provided critical revisions and contributed to theoretical development.

## Acknowledgments

The authors would like to thank Dr. Jun Noguchi and Dr. Satoshi Watanabe for encouraging discussions. Thanks also to Akiko Tsuchiya for caring marmosets, and special thanks to all the subject marmosets.

