## Supplemental Figure 1 & 2 for "Hyperactivity is linked to elevated cortisol levels: comprehensive behavioral analysis in the prenatal valproic acid-induced marmoset model of autism"

### Supplemental materials

**Figure S1**: Average body weights (in grams) and age (weeks) of the adult subjects in the home-cage activity analysis. No significant between-group differences were detected (weight: $p = 0.9634$; age: $p = 0.8541$ for Student's *t* test).

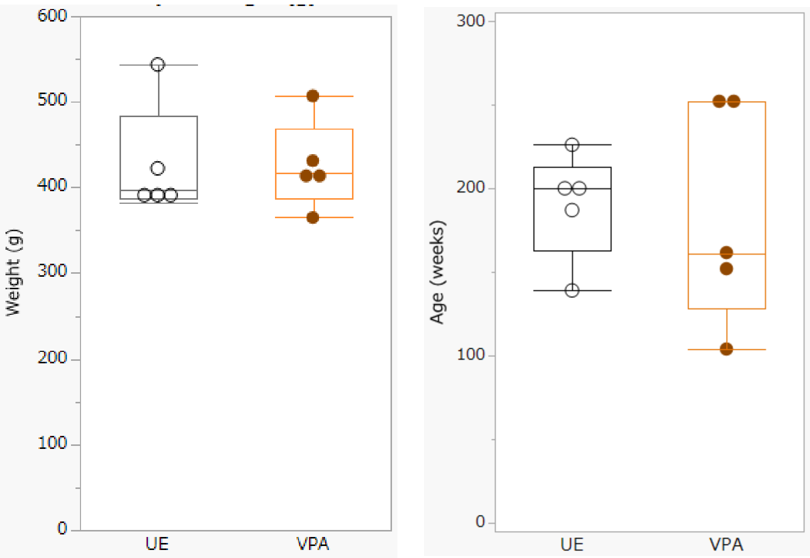


**Figure S2**: Average weights (in grams) of the juvenile subjects in the 3-chamber test. There was no significant difference in weights between the two groups ($p = 0.0878$ for Student's *t* test).

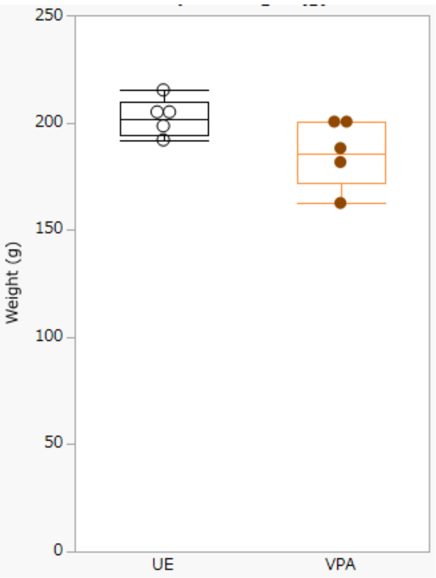
